# Leveraging eDNA metabarcoding to characterize nearshore fish communities in Southeast Alaska: Do habitat and tide matter?

**DOI:** 10.1101/2021.10.28.466160

**Authors:** Wes Larson, Patrick Barry, Willie Dokai, Jacek Maselko, John Olson, Diana Baetscher

**Author notes:** Corresponding author: Wesley A. Larson).

## Abstract

Nearshore marine habitats are critical for a variety of commercially important fish species, but assessing fish communities in these habitats is costly and time-intensive. Here, we leverage eDNA metabarcoding to characterize nearshore fish communities near Juneau, Alaska, USA, a high-latitude environment with large tidal swings, strong currents, and significant freshwater input. We investigated whether species richness and community composition differed across three habitat types (sand beaches, eelgrass beds, and rocky shorelines) and between high and low tides. Additionally, we tested whether replication of field samples and PCR reactions influenced either species richness or composition. We amplified a 12S mitochondrial locus in our samples and identified 188 fish amplicon sequence variants (ASVs), corresponding to 21 unique taxa, with approximately half of these resolved to single species. Species richness and composition inferred from eDNA differed substantially among habitats, with rock habitats containing fewer taxa and fewer overall detections than sand and eelgrass habitats. The effect of tide was more subtle and suggested a habitat-tide interaction, with differences in taxa between tides largely isolated to sand habitats. Power analyses indicated that additional field sampling is useful to detect subtle changes in species richness such as those due to tide. PCR replicates typically identified a small number of additional taxa. The most notable result from our study was that shore morphology appeared to substantially influence community structure. Rocky shorelines sloped quickly into deep water, while sand and eelgrass habitats descended much more gradually. We hypothesize that differences in taxa observed among habitats were largely due to lack of mixing between bottom and surface water, providing further evidence that eDNA transport is minimal and that many marine eDNA detections are derived from highly localized sampling locations. We suggest that future studies could explore the extent to which habitat and nearshore physical processes influence eDNA detections.

## Introduction

Understanding the distribution of fish species across space and time is vital for the development of conservation and management strategies to protect these species and promote productive and sustainable fisheries (Baudron et al., 2020; Thorson & Barnett, 2017). Assessing fish communities is especially important in nearshore marine habitats, as these areas are critical for a variety of commercially valuable species, particularly in their sensitive early life stages (Craig et al., 2006). Traditionally, many surveys of fish communities in nearshore habitats have been conducted using beach seines or similar gear types deployed from either shore or small boats (Grüss et al., 2021; Laurel & Rogers, 2020). However, these methods are time and resource intensive, requiring substantial infrastructure and personnel time (Steele, Schroeter, & Pace, 2006). Alternatively, sampling and then analyzing environmental DNA (eDNA) can generate similar datasets with substantially less effort (Shelton et al., 2019).

Pioneering research with marine eDNA demonstrated that eDNA metabarcoding recovered fish diversity as well or better than conventional methods within a small harbor in Denmark (Thomsen et al., 2012). More recently, single and multi-gene approaches have been used to address ecological questions such as community changes across time (Djurhuus et al., 2020), interannual changes in oceanographic conditions (Closek et al., 2019), and fine-scale community variation (West et al., 2020). Importantly, previous studies have used marine eDNA metabarcoding to detect differences in fish communities across habitats at small spatial scales, which is vital for developing habitat conservation strategies (Sigsgaard et al., 2020). Differences in fish communities were detected at scales ∼60 m across a 2.5 km transect spanning multiple diverse habitats (Port et al., 2016), at <100 m (O’Donnell et al., 2017), and across 140 m between the nearshore and surf zone associated with 4-5 m changes in depth (Monuki, Barber, & Gold, 2021).

Despite the rapid adoption of eDNA as a method for sampling marine environments, studies of mechanisms that can influence the distribution of eDNA have lagged behind, in part because many mechanisms are localized and require empirical studies spread across habitats and latitudes. Although eDNA studies have detected fine-scale differences in fish communities across habitats and depths, water movement from tidal exchange and currents could redistribute eDNA away from the organisms from which the DNA originates. At least one study that used regional ocean modeling suggested that eDNA in the coastal ocean could be transported tens of kilometers in a few days (Andruszkiewicz et al., 2019). However, the effects of tides and currents are geographically constrained. The only eDNA study that investigated the influence of tides found that species communities were similar across a ∼3 m tidal range (Kelly, Gallego, & Jacobs-Palmer, 2018). Specifically, Kelly et al. (2018) sampled eDNA from three sites in a natural glacial fjord across a two-day period on incoming and outgoing tides and found no substantial differences in community structure that could be attributed to tide.

The distribution of eDNA across nearshore habitats is influenced by water movement as well as DNA degradation rate (Harrison, Sunday, & Rogers, 2019). Marine eDNA is degraded by a combination of chemical, physical, and biological factors (Holman, Chng, & Rius, 2021), and although eDNA from different species has been found to decay at different rates under the same environmental conditions, multiple studies identify changes in eDNA detections after 48 hours (Collins et al., 2018; Holman et al., 2021).

Given the potential for eDNA to be influenced by large-scale water movements, it is surprising that so many previous studies detect fine-scale heterogeneity and demonstrate little influence of tide. This disconnect highlights the dramatic variation across nearshore habitats and suggests that additional studies are necessary to understand the variables that influence community composition data derived from marine eDNA.

High latitudes warrant additional work to understand mechanisms influencing eDNA because these cold, productive waters support some of the most important fisheries in the world and are often influenced by strong currents, high tidal swings, and substantial storm activity (Skern-Mauritzen, Olsen, & Huse, 2018; Weingartner, Eisner, Eckert, & Danielson, 2009). One high-latitude area where eDNA studies could be highly beneficial for assessing species diversity is within Alaska waters. Alaska supports some of the most valuable fisheries in the world and includes 6,640 miles (10,686 km) of coastline (Beaudreau et al., 2019). However, Alaska’s vast geography with diverse coastal habitats makes conducting representative sampling logistically challenging. These challenges can result in fisheries being managed based on limited data, particularly in nearshore habitats (Newman, Berkson, & Suatoni, 2015). eDNA represents a powerful method to rapidly increase data availability for diverse species across Alaska (see Liu et al., 2019; Stoeckle et al., 2020 for examples outside Alaska), but to date no studies using eDNA metabarcoding to characterize marine fish communities have been conducted in this region.

Here, we conduct the first marine eDNA metabarcoding study focused on fish communities in Alaska with the goal of validating this method in a high-latitude nearshore marine environment. We sampled eDNA from nine sites spanning a large habitat gradient near Juneau in Southeast Alaska (Fig. 1), a region characterized by high tidal swings, strong currents, and significant freshwater input from rivers and rainfall. Our specific objectives were to (1) investigate potential differences in species richness and fish community composition among habitat types, (2) evaluate the effect of sampling at different tidal stages, and (3) investigate the influence of sample replication on inferences of species richness. This study provides important information on factors influencing estimates of community composition derived from eDNA that can be generalized beyond this study and also presents a standardized workflow that can serve as the foundation for future eDNA sampling in Alaska.

**Figure 1:**
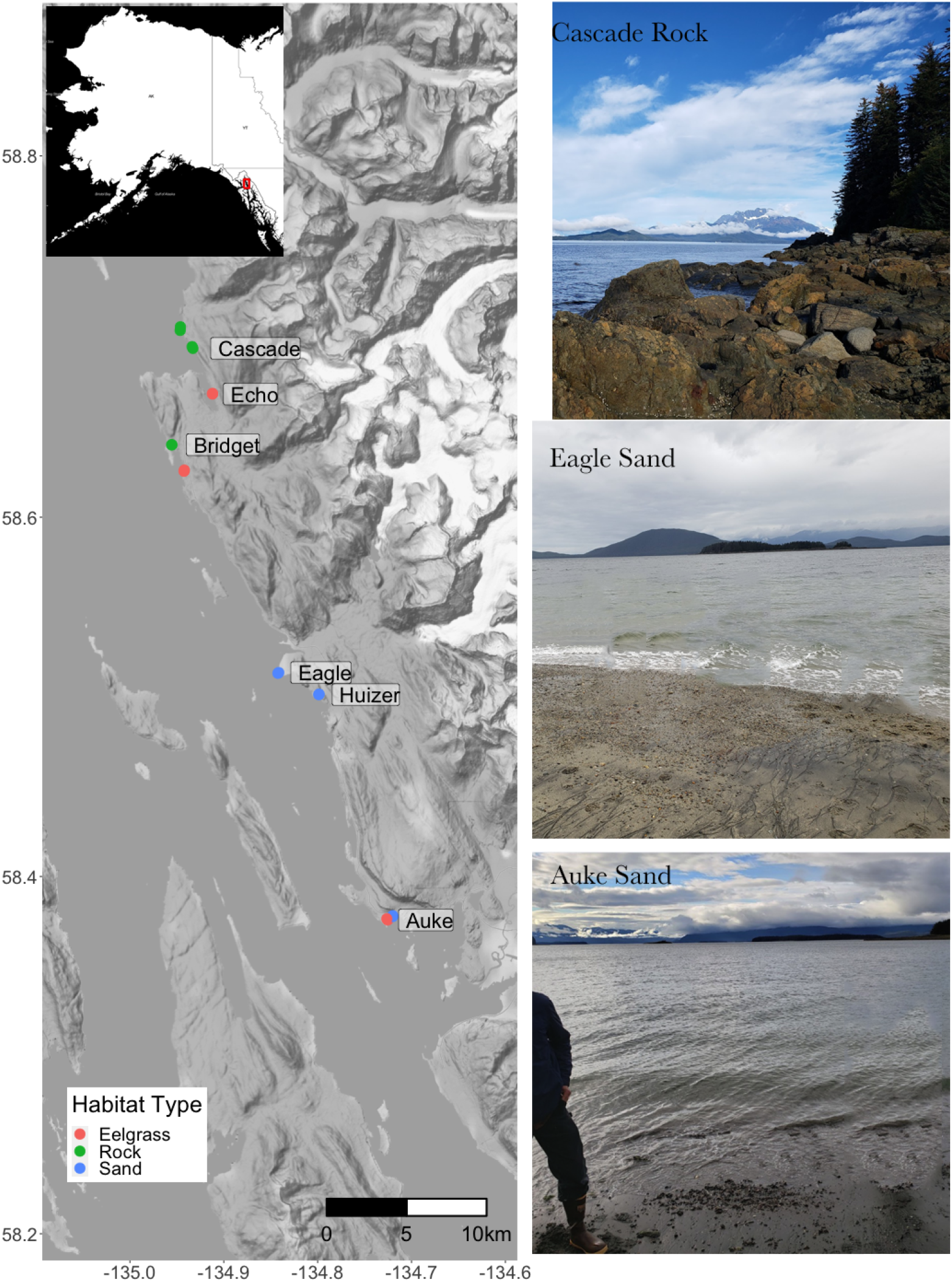
Map of the nine nearshore sampling locations around Juneau, Alaska, sampled for eDNA in early fall of 2020. Pictures illustrate the typical sand and rock habitats encountered during sampling. Eelgrass habitats are composed of eelgrass growing on a sandy substrate and are visually similar to sand. See Tables 1 and S1 for more information on sampling sites and see the Methods section for more detailed habitat descriptions.

## Methods

### Study location and sample collection

We sampled eDNA from shore at nine sites near Juneau in Southeast Alaska, USA, during a single week in early fall of 2020 (Fig. 1, Table 1). These sites were all within 50 km of each other and were chosen based on data from the Nearshore Fish Atlas (Johnson, Neff, Thedinga, Lindeberg, & Maselko, 2012), which integrates habitat and biological survey data from multiple sources. Sites were chosen to encompass the major habitat types in the region: sand/gravel, kelp/bedrock, and eelgrass. Sand/gravel beaches (hereafter referred to as sand) are composed largely of sand and small gravel with few larger rocks or macroalgae, kelp/bedrock areas (hereafter referred to as rock) are composed of large boulders and bedrock which can be seasonally (spring to summer) covered in large macroalgae such as bull kelp (*Nereocystis luetkeana*) but were not during the study period, and eelgrass (*Zostera marina)* beds, which are important habitat for small fish and generally found near sandy beaches (Hogrefe, Ward, Donnelly, & Dau, 2014). Sand and eelgrass environments are often gradually sloping, with large sections of beach exposed at low tide, whereas rock environments are often characterized by abrupt drop-offs into waters > 10 m deep.

**Table 1:**
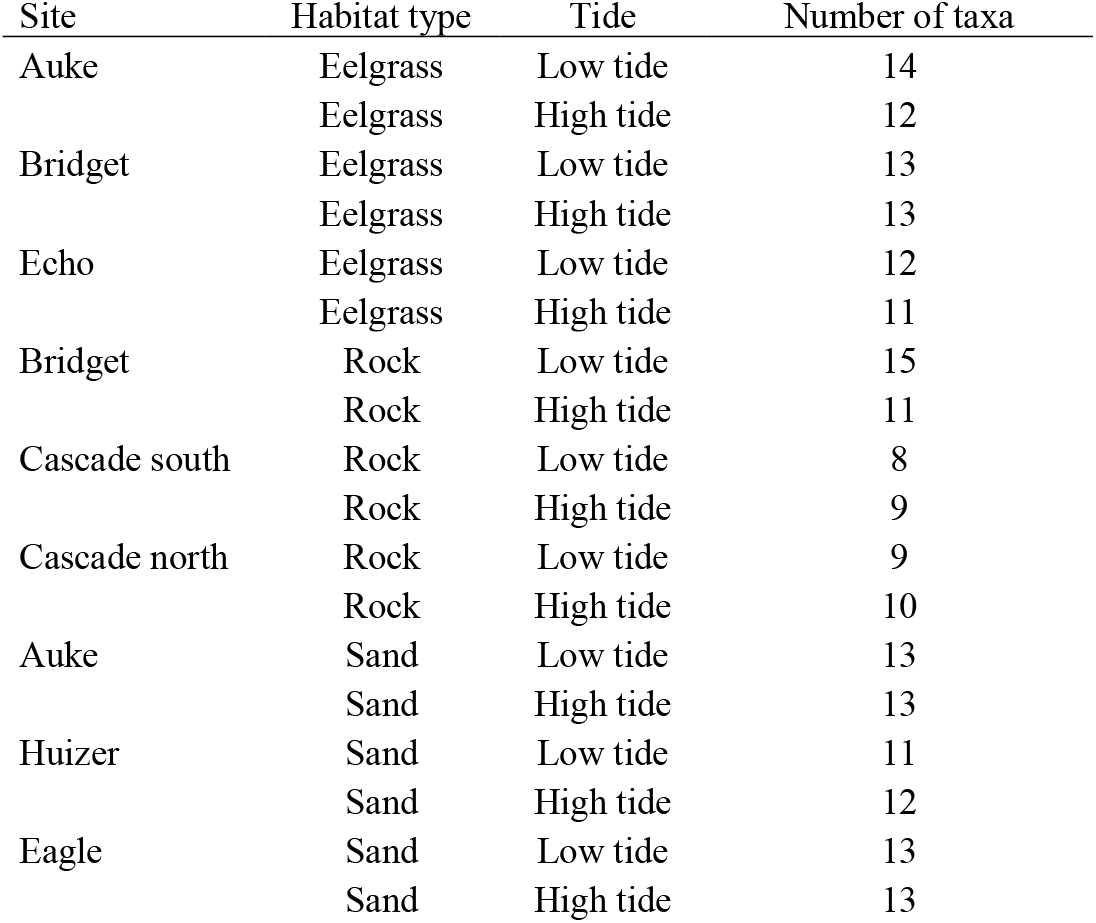
Sampling sites, habitat type, and the number of unique taxa detected at each site during high and low tides. See Table S1 and Fig. 1 for additional information about sampling sites. A detailed description of each habitat type is found in the Methods section.

We collected nine 1-L water samples and a negative control from each of the nine sites during a single high and low tide cycle (Juneau has mixed semidiurnal tides). All samples were taken within 30 minutes of slack tide, the time of minimum tidal water movement (Table S2). Sand and eelgrass sites were sampled by hand just below the surface in approximately 1 m of water using waders, while rock sites were sampled just below the surface from shore without entering the water. The nine replicate samples for each site/tide combination were collected randomly in an approximately 100 m^2^ area. High- and low-tide samples from each site were collected on the same day to minimize temporal changes in species communities not caused by tide, and samples were taken between 8 am and 8 pm to minimize diurnal species community changes.

Collection, filtration, and decontamination methods generally followed those of Gehri, Larson, Gruenthal, Sard, and Shi (2021). Each 1-L Nalgene collection bottle was decontaminated for 10 minutes in 20% chlorine bleach solution and then rinsed at least three times with deionized water. Field negatives consisted of 1-L bottles filled with laboratory grade water, brought to the field, opened for 30 seconds during sample collection, and subsequently handled like all other samples. All samples were filtered through new and sealed 47 mm diameter, 0.45 µm pore nitrocellulose Nalgene analytical test filter funnels. Non-sterile filtering supplies and instruments were sterilized by soaking in a 20% chlorine bleach solution for at least 10 min and then rinsed at least 3 times with deionized water. Filters were preserved in ∼95% ethanol in 2 ml tubes and stored at room temperature.

### Sequencing library preparation

DNA was extracted within two months of collection following Gehri et al. (2021) with Qiagen DNeasy Blood & Tissue Kits and the Purification of Total DNA from Animal Tissues Spin-Column Protocol (Thermo Fisher). Extractions were performed in a UV light sterilized, HEPA filtered, laminar air flow hood system, in which surfaces were frequently treated with 20% bleach solution to reduce the risk of contamination. Eluted DNA was treated with Zymo OneStep PCR Inhibitor Removal columns (Zymo Research).

PCR was performed on extracted DNA with primers that amplify a ∼142 bp region of the 12S mitochondrial genome (Riaz et al., 2011). PCR reactions were performed in 10 μl volumes using 3 μl of template eDNA, and 7 μl of PCR master mix. The PCR master mix consisted of per reaction volumes of: 1 μl of New England Biolabs 10X Standard Taq Reaction Buffer, 0.2 μl of 10 mM dNTPs, 0.8 μl of 25 mM MgCl2, 0.5 μl of 20 mg/ml bovine serum albumin (BSA), 0.3 μl of 1.25 U/µl NEB Taq, 2.6 μl of molecular grade water, and 0.8 μl of 10 μM forward and reverse primer. Thermal cycling was performed as follows: 95°C for 2 min, followed by 35 cycles of 95°C for 30 s, 57°C for 30 s, 72°C for 45 s, and then a single 72°C extension for 5 min.

One PCR negative control, four extraction negative controls, and a positive control were included on each 96-well PCR plate. PCR negative controls consisted of 3 μl of laboratory grade water. Extraction negative controls consisted of an unused Nalgene filter that was extracted alongside, and in the same manner as, eDNA samples. We used walleye (*Sander vitreus)*, a freshwater fish from the midwestern USA and Canada that is not present in Alaska, as a positive control. Finally, to estimate variation in species detection across DNA aliquots from the same extraction, we included eight PCR replicates for 11 samples and an additional four PCR negative controls, and four positive controls.

Sequencing libraries were prepared using the Genotyping-in-Thousands by Sequencing (GT-seq) protocol (Campbell, Harmon, & Narum, 2015). PCR products were indexed in a barcoding PCR, normalized using SequalPrep plates (Invitrogen) and each 96 well plate was subsequently pooled. Next, a double-sided bead size selection was performed using AMPure XP beads (Beckman Coulter), using ratios of beads to library of 0.5x to remove non-target larger fragments and then 1.2x to retain the desired amplicon. Libraries were sequenced on a MiSeq (Illumina) using a single 150-cycle lane run with 2×75 bp paired-end (PE) chemistry.

### Data filtering and quality control

Methods for data filtering and quality control were similar to those described in Gehri et al. (2021). PE reads for each individual were joined with FLASH2 (Magoč & Salzberg, 2011; https://github.com/dstreett/FLASH2) with default settings except for the maximum overlap was set to 10 (based on an amplicon length of 142bp and 2×75bp chemistry). We then used the program DADA2 (Callahan et al., 2016) to quality filter reads (specifying a total length of 142bp and a maximum number of expected errors = 2), remove chimeras (with the consensus method), and export amplicon sequence variants (ASVs) and the number of times they occurred in each sample. ASVs were compared to all sequences in the National Center for Biotechnology Information (NCBI) nucleotide database using BLASTn (Altschul, Gish, Miller, Myers, & Lipman, 1990). We then parsed aligned ASVs with custom R scripts (available from https://github.com/AFSC-Genetics/eDNA_NearshoreMarine, repo to be made public upon acceptance) and retained matches with greater than or equal to 98% sequence identity and alignment lengths greater than 139bp (similar parameters to Gehri et al., 2021 who amplified the same gene). At this stage, we removed all ASVs that did not align to fish or that did not contain a match that passed our minimum thresholds. ASVs with a single match that met our parameters or that had multiple matches to a single species (unambiguous) were assigned to that species. However, many ASVs matched multiple species with sequence similarity greater than our 98% threshold. In these cases, we iteratively increased taxonomic levels (i.e., genus, family, order) until matches were unambiguous. We also used species distribution information from FishBase (http://www.fishbase.org/) to exclude matches that were outside of the geographic range of our study area (i.e., Northeast Pacific) but genetically similar to species found within it. ASVs assigned to the same taxonomic unit were collapsed to form a dataset containing the number of reads for each taxonomically distinct ASV in each sample.

To control for contamination, from each water sample we subtracted the maximum number of reads of each ASV detected in the negative and positive controls. We then transformed this read count data into binary presence/absence data, with ASVs having > 4 reads in a given sample considered a positive detection.

### Statistical analysis

We used a variety of data visualization techniques, univariate, and multivariate analyses to investigate whether habitat and tide influenced species richness and community composition inferred from our eDNA samples. For species richness analysis, the number of taxa detected was computed for each sample, whereas for community composition analysis, data were coded as the number of samples with positive detections for a given taxon at a given site, with 9 being the maximum number of detections possible per site. First, to visually assess differences in species detections across different habitats and tide stages, we generated a heatmap of the number of detections of each ASV at each site and tide stage. We also calculated average species (taxon) richness by habitat and tide stage as well as taxon-specific differences in detection rates among habitats and tide stages.

Next, to assess the influence of tide and habitat on community composition, we conducted distance-based redundancy analysis (db-RDA) based on Bray-Curtis dissimilarity in the R package vegan (Oksanen et al., 2020). The db-RDA hypothesis testing framework allows testing for habitat and tide interaction effects (Legendre & Anderson, 1999). The significance of each environmental term (i.e., their influence on variation in community composition) was assessed with ANOVA-like permutation tests.

To test for the effects of tide and habitat on species richness, we used a generalized linear mixed-effects model (GLMM; alpha = 0.05) and the package lme4 (Bates, Mächler, Bolker, & Walker, 2015), with sampling replicates within the same habitats treated as random effects. The GLMM error distribution family was determined using the poissonness plot method (Hoaglin, 1980) and distplot function in the vcd R package (Meyer, Zeileis, & Hornik, 2006). We tested the Poisson, binomial, and negative binomial family distributions for best fit. We also conducted a power analysis to determine the effect of the number of replicate samples per site required to detect differences in species richness associated with tide and habitat. We did this by drawing between 3-9 samples with replacement from each site and conducting the GLMM analysis as described above. After 1,000 bootstrapped iterations of our simulated sampling, we calculated the probability of detecting a significant effect (alpha = 0.05) for each sample size. This method allowed us to determine whether fewer replicate samples per site would have been sufficient to detect the trends observed.

Finally, we tested whether additional taxa were detected by performing nine PCR replicates for a subset of 11 samples. We used species accumulation curves to determine how many replicates were required to obtain the maximum number of taxa for a given sample (i.e., the taxon was detected in at least one of 9 PCR replicates with > 4 reads).

## Results

### Sequencing, taxonomic assignment, and quality control

We obtained a total of 11,033,262 PE reads from 250 samples with an average of 44,134 reads per sample. After PE assembly, filtering, merging, and chimera removal in DADA2, 969,988 reads remained (17.6%). These reads corresponded to 307 ASVs, 9 of which failed to align to a sequence within NCBI, 49 were filtered out because they matched sequences with low similarity (<98%) or were short sequences (<140 bp), and 61 ASVs matched non-fish organisms. We retained 21 unique taxa after collapsing ASVs into taxonomic groups that could be distinguished reliably based on sequence variation, one of which was found in only a single PCR replicate (*Aulorhynchus flavidus*). Some taxa consisted of single species that were highly distinguishable (e.g., *Artedius fenestralis* [padded sculpin], *Oxylebius pictus* [painted greenling], *Ophiodon elongatus* [lingcod]), whereas it was necessary to group other taxa into higher taxonomic levels such as genus (Hexagrammos), family (e.g., Gasterosteidae), or infraorder (e.g., cottales).

Contamination that occurred during sampling in the field, PCR, and extraction was generally low, with fewer than 10 reads per taxon except in a few cases. However, we did detect >1,000 reads for Salmoninae and Pleuronectidae in a two field negative samples (Table S2). Both PCR and extraction negative controls had minimal contamination (<50 reads for a single taxa), indicating that the small amounts of contamination in our study were likely introduced either in the field or during the filtering process. Over 99.99% of the reads in positive controls assigned to walleye, while reads from this species were rare in other samples (average proportion of walleye reads in water samples was 4.9% and 78% of the samples had <5% of reads from walleye).

### Influence of habitat and tide

Taxa identified in our study can be divided into three primary categories: (1) common taxa detected at all or nearly all sites, (2) taxa detected at moderate levels in some but not all sites, and (3) rarely detected taxa (Fig. 2). The most common taxon was Salmoninae (salmon); this taxon was detected at all sites and tides. Other common taxa included *Clupea pallasii* (Pacific herring), Pleuronectidae (flatfish), Zoarcales (pricklebacks and gunnels), Cottales (sculpin), and *Leptocottus armatus* (Pacific staghorn sculpin). Taxa that were detected at some but not all sites included Gadidae (cods), Anoplopomatidae (sablefish), and Osmeriformes (smelt). Notable rarer taxa include Sebastinae (rockfish), and *Ophiodon elongatus* (lingcod).

**Figure 2:**
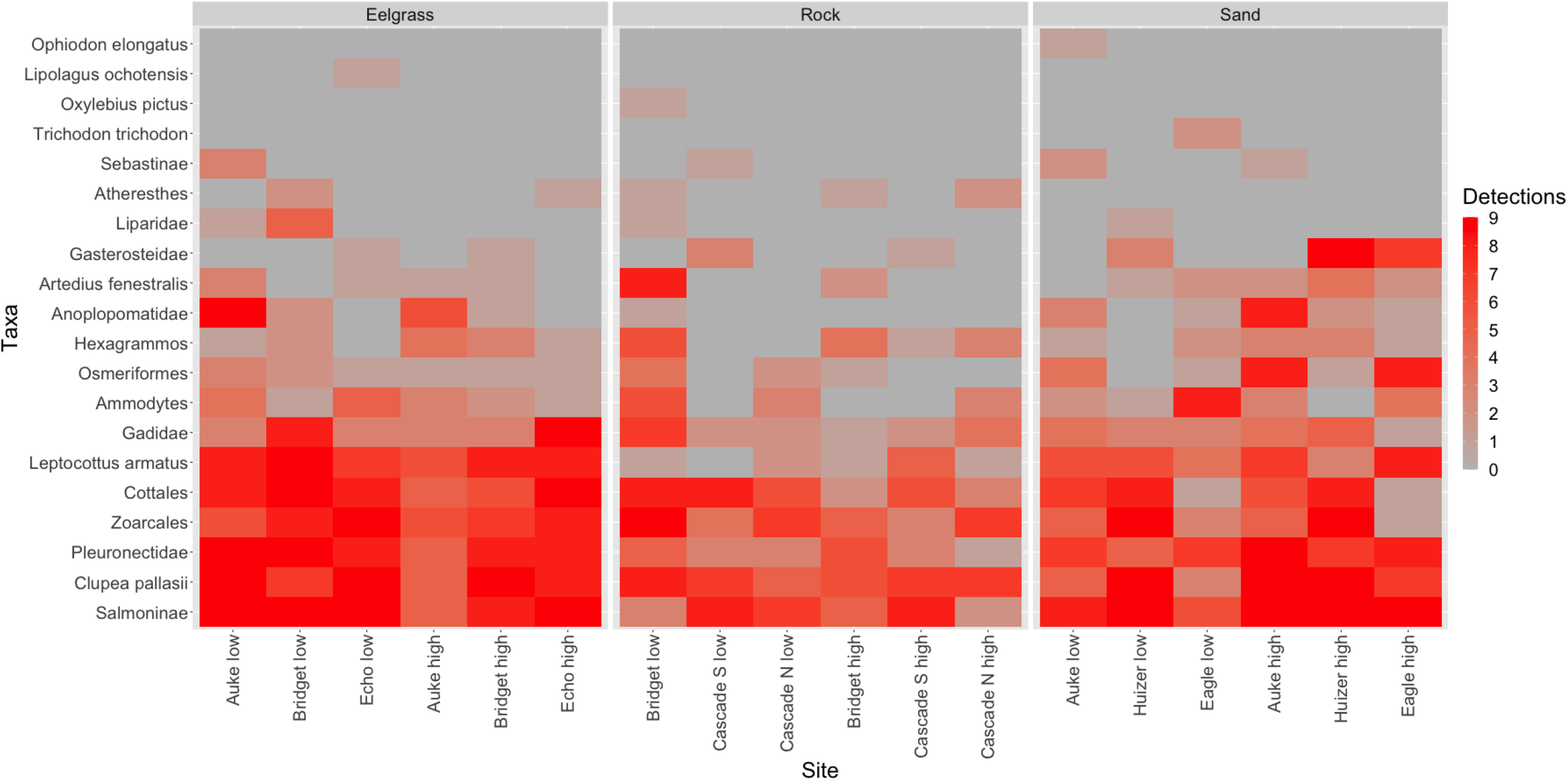
Heatmap of the number of detections (out of 9 samples) for each taxon and sampling site.

On average 12 taxa were detected per sampling event, with the highest number (15) detected at the Bridget rock low tide site and the lowest (8) at the Cascade south rock low tide site (Table 1). The number of taxa detected differed by habitat, with 13 taxa on average detected at sand and eelgrass sites and 10 detected at rock sites. The number of taxa detected at high and low tide sites was the same with an average of 12.

In general, rock habitats contained fewer species, different species, and fewer positive detections for each species compared to sand and eelgrass habitats (Tables 1, 2; Figs 2–4). However, the influence of tide was more subtle (Table 1,2; Figs 2,3,5). The permutation tests based on db-RDA with a tide and habitat interaction showed a significant habitat effect (*p* = 0.004), but no significant effect for tide (*p* = 0.428), or tide and habitat interaction (*p* = 0.723). Visualization of db-RDA results without the non-significant interaction variable demonstrated a clear separation of rock sites from both sand and eelgrass sites, but no apparent clustering by tide (Fig. 3). The habitat rock variable displayed high loadings on RDA1, which explained 20% of the variation, while the habitat sand and tide variable displayed high loadings on RDA2, which explained only 6% of the variation when the non-significant interaction term was removed.

**Table 2:** Analysis of deviance from generalized linear mixed-effects model investigating the influence of tide and habitat on species richness.

| Terms | $X^2$ | DF | $p$ value |
| --- | --- | --- | --- |
| Tide | 0.97 | 1 | 0.32 |
| Habitat | 20.92 | 2 | 0.00003 |
| Tide:Habitat | 15.83 | 2 | 0.0004 |

**Figure 3.**
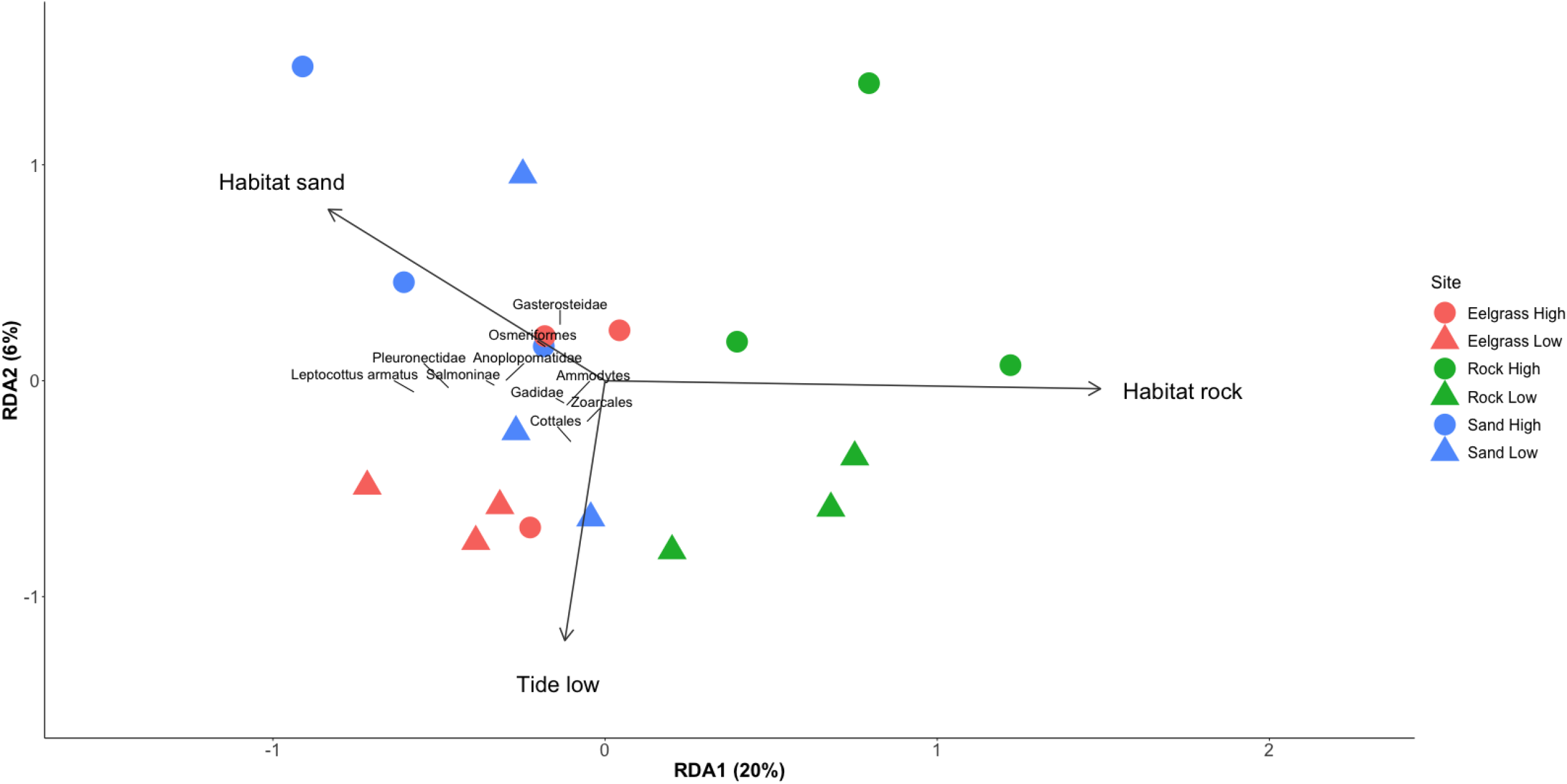
Distance based redundancy analysis (db-RDA) illustrating the influence of habitat and tide on species composition across nearshore marine sites. Numbers represent sites in Table 1. Only the 10 taxa with the highest scores (grey text) were included in the plot to facilitate visualization. In the permutation tests, the habitat term was significant (*p* = 0.004) while tide and interaction terms were not (*p* = 0.429 and 0.722 respectively).

**Figure 4:**
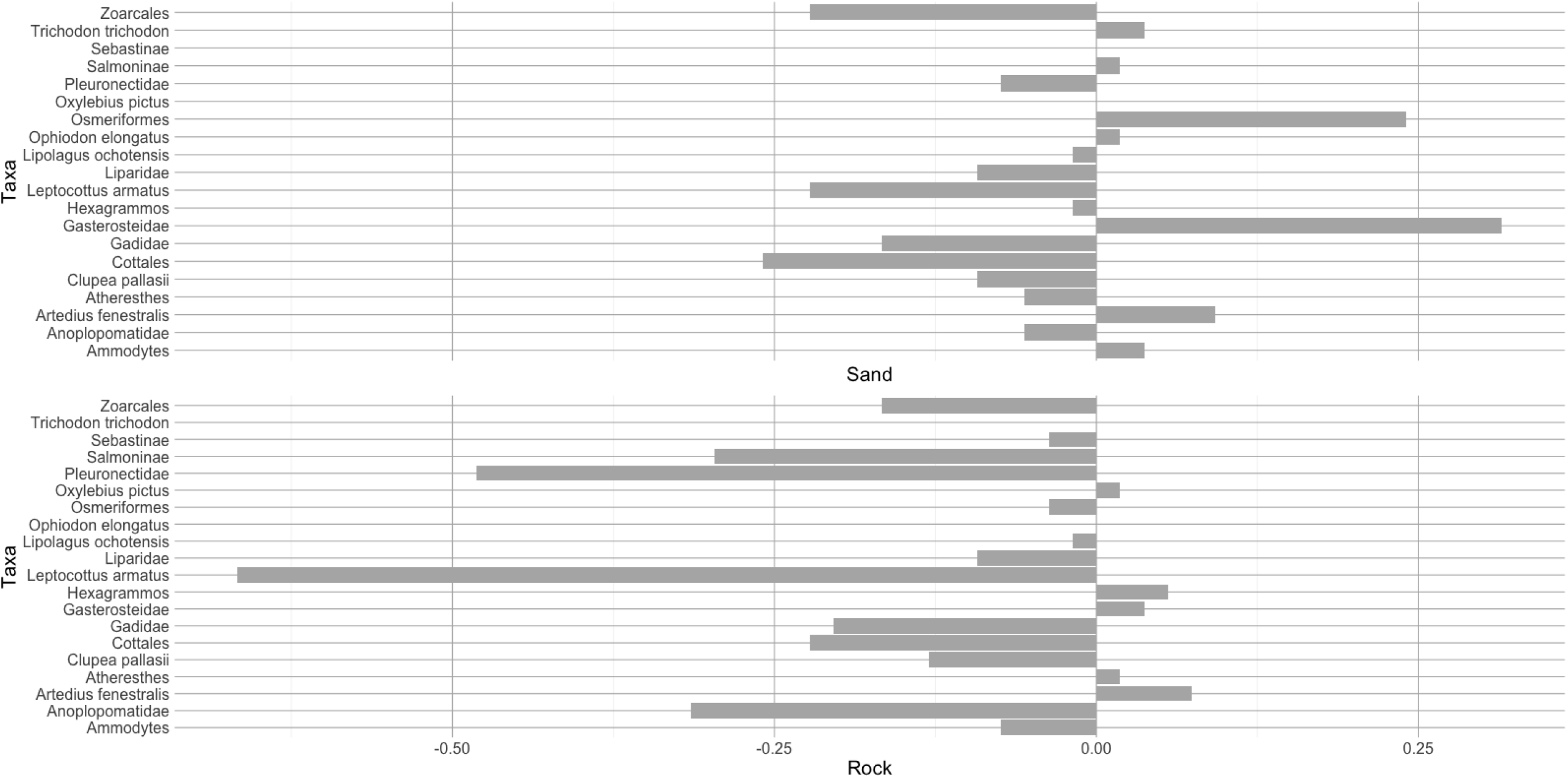
Change in the detection rate of each taxon in sand and rock habitat as compared to eelgrass (the habitat type with the greatest number of taxa) across both tide stages. Positive values indicate that the taxon was detected more often in sand or gravel compared to eelgrass and negative values indicate that the taxon was detected less often.

**Figure 5:**
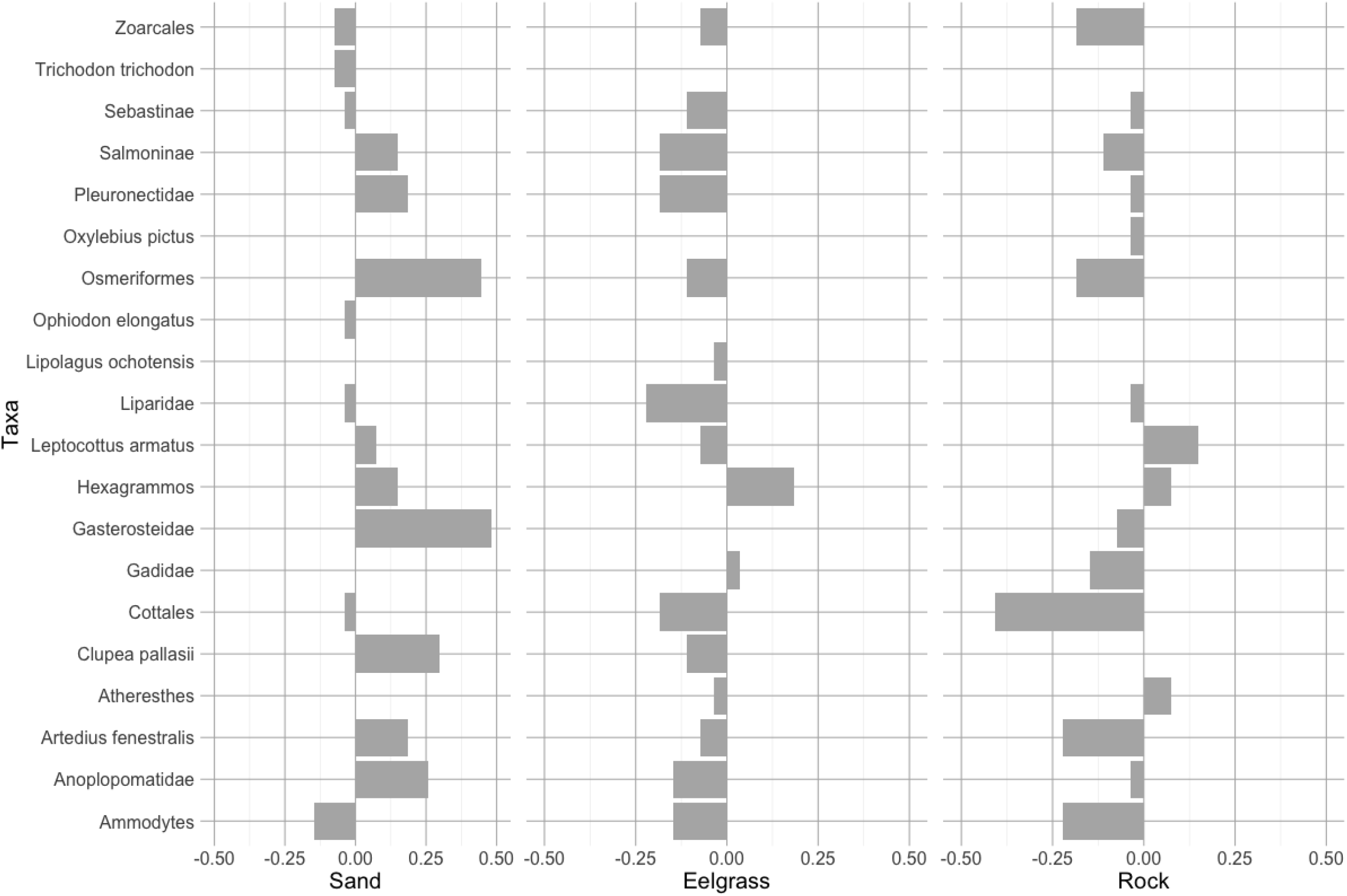
Differences in detection rate of each taxon by tide for each habitat type. Positive values indicate that the taxon was detected more often at high tide and negative values indicate that the taxon was detected more often at low tide.

Results from the mixed-effects model were concordant with the RDA, with high support for an effect of habitat on species richness (P < 0.001), but little support for an effect of tide (*p* = 0.32; Table 2). However, the mixed-effects model did indicate support for an interaction effect between habitat and tide (*p* < 0.001), suggesting that tide may influence species richness differently depending on habitat.

The primary taxa driving differences between habitats were *Leptocottus armatus*, Pleuronectidae, Anoplopomatidae, and Salmoninae, with all of these taxa being less common in rock habitats (Figs. 2-4). Taxon composition was relatively similar across tides except for in sand habitats, where small pelagic fish were more commonly detected at high tide (Fig. 5). Specifically, Gasterosteidae, Osmeriformes, and *Clupea pallasii* were all detected much more frequently at high versus low tides. Tide appeared to influence species compositions substantially in sand but not the other habitats.

### Investigating the effects of replication

The bootstrapped GLMM mixed-effects power analysis showed the larger effect of habitat than tide on species composition. Three samples would have been sufficient to detect significant effects of habitat (probability of detection > 0.7), with significant effects detected more consistently with an increasing number of replicates (probability of detection > 0.9 with nine replicates). However, finding a significant tide-habitat interaction effect appeared dependent on adequate sampling, requiring four replicates to obtain a probability of detection greater than 0.5 and seven replicates to obtain a probability greater than 0.9 (Table 3). Tide was rarely found to be significant (probability of detection ≤ 0.1) suggesting that the effect size of tide was too small to detect with nine replicate samples or that there was no difference between samples taken at different tides (Table 3).

**Table 3:** Results of generalized linear mixed-effects models conducted on bootstrapped datasets (1,000 iterations) with 3-9 samples selected with replacement per sampling event. The N samples column is the number of samples included in the bootstrapped datasets, and the other columns are the proportion of bootstrap replicates where the model produced a significant *p* value (*p* < 0.05) for a given term (i.e. power of the test), which corresponds to the probability of detection. For example, nearly all bootstrap replicates (0.972) were significant for the tide-habitat interaction with nine samples.

| N samples | Tide | Habitat | Tide-Habitat |
| --- | --- | --- | --- |
| 3 | 0.036 | 0.729 | 0.482 |
| 4 | 0.046 | 0.84 | 0.635 |
| 5 | 0.072 | 0.863 | 0.781 |
| 6 | 0.080 | 0.876 | 0.875 |
| 7 | 0.066 | 0.908 | 0.914 |
| 8 | 0.105 | 0.894 | 0.946 |
| 9 | 0.100 | 0.905 | 0.972 |

Additional taxa were detected in some of the PCR replicates for the subset of samples tested. Six of 11 samples identified all taxa in either a single PCR replicate or cumulatively between two replicates, and five replicates were required for cumulatively detecting all taxa in only one of the 11 samples (Fig. S1). Common taxa, such as Salmoninae and Zoarcales, were almost always detected in > 75% of replicates for samples in which they were present. However, detections of rarer taxa were more sporadic across PCR replicates. For example, Gadidae was often detected in < 25% of replicates. Finally, we detected one additional taxon (*Aulorhynchus flavidus*, tube-snout) in PCR replicates that was not detected in the rest of the study; this detection occurred in the Auke eelgrass low tide sample.

## Discussion

### Influence of habitat and tide on community composition

Habitat had a significant effect on fish species richness and community composition inferred from eDNA, while the influence of tide was less obvious. In particular, fish communities differed significantly between rock habitats and the other two habitats that we sampled, sand and eelgrass. Tide also influenced fish community composition, although the impact of tide was more subtle and habitat dependent, with sand being the only habitat displaying substantial differences in the fish community between tides.

eDNA samples taken in rock habitats may be subject to greater water stratification than sand or eelgrass habitats. Bottom depths at sampling locations in rock habitats were often deeper, with slopes that descend rapidly from the collection location, sometimes reaching 100 m within just a few meters distance from shore. In contrast, sand and eelgrass environments are characterized by more gradual slopes. Since many of the taxa that we detected are strongly associated with the benthos (e.g., Pleuronectidae and *Leptocottus armatus*), it is possible that the lower detection rates for these species in rock habitats are a function of sampling farther from the benthic habitat in locations where water is not well-mixed from surface to benthos. Additionally, the increased vertical distance (depth) from the benthos to the surface associated with the rock habitats may have diluted eDNA from benthic species compared to the shallower water column and shorter sampling distance from surface to benthos in sand and eelgrass habitats. Additionally, we speculate that in rock habitats, the halocline may act as a barrier to eDNA movement, effectively trapping surface and benthic sourced eDNA in their respective layers. The halocline may be especially pronounced in coastal southeast Alaska due to large freshwater inputs from rivers, glaciers and precipitation. Our sampling protocol did not collect information about the temperature or salinity of the water mass sampled, but deploying a CTD during future sampling would provide more information to test this hypothesis.

Multiple eDNA studies have identified differences in community composition in samples taken at different depths (Andruszkiewicz et al., 2017; Monuki et al., 2021) or water layers (Littlefair, Hrenchuk, Blanchfield, Rennie, & Cristescu, 2021). In particular, a recent study detected extremely fine-scale differences in community composition inferred from eDNA at scales of less than 3 m in a kelp forest in California, USA (Monuki et al., 2021). This and other empirical studies, including ours, provide evidence that eDNA transport is likely more limited than suggested by previous modeling studies (e.g., Andruszkiewicz et al., 2019). However, our study system differs substantially from the complex and highly structured kelp forest habitats that have illustrated fine-scale heterogeneity in nearshore eDNA results (Monuki et al., 2021; Port et al., 2016). Instead, the general species composition in our study system appears to be similar across habitats, but the detection rates of these species appear to be influenced by habitat features (i.e., slope, depth). Further, our study highlights how the species community derived from eDNA can be influenced by depth even when samples are taken at the same depth (surface) and at approximately the same distance from shore.

The substantial tidal exchanges near Juneau, Alaska, suggest that water masses would be mixed and replaced frequently. However, our results provide evidence that eDNA transport may be limited, despite this daily water movement. Monuki et al. (2021) also provided evidence that eDNA transport may be limited in a dynamic system with much greater wave action than is present in the nearshore environments sampled in our study. They suggest that eDNA from local sources may dominate due to high and continuous shedding rates from local individuals. It seems likely that the dilution of eDNA in the ocean coupled with the frequent shedding of eDNA from local individuals leads to a dominant signature of local sources in eDNA samples.

Although we found a large effect of habitat on fish community composition, the effect of tide was much more subtle. Differences between tides were primarily only found in sand habitats and were largely driven by higher detection rates of small pelagic fish such as Gasterosteidae (sticklebacks), Osmeriformes (smelts) and *Clupea pallasii* (Pacific herring) at high tide. This trend was especially apparent in the Eagle Beach samples. Eagle beach differs from other beaches in our study because it is part of a large glacial alluvial fan formed by the Eagle/Herbert River complex. This beach is characterized by different bathymetry than other sites as well as a large freshwater influx. We hypothesize that the increased detections of small pelagic fish at high tide were a reflection of movement of these fish to either (1) seek shelter from predators, (2) exploit a food source, or (3) other tide-associated movement into freshwater.

The only previous study that explicitly tested the effects of tide found no consistent differences between tidal cycles in the Puget Sound, Washington, and concluded that sampling location influenced eDNA results more substantially than tidal stage (Kelly et al., 2018). Our results, that habitats affect fish community more than tides, generally match this conclusion but also highlight the ability for eDNA to detect temporary movements of vagile organisms in response to different tidal stages. Taken together, the results of our study and Kelly et al. (2018) suggest that the majority of species in the fish community will be detected with eDNA regardless of tidal stage but that habitat or species-specific differences may exist between tides.

### Replication and optimization of sampling effort

Optimizing sampling effort is a key logistical consideration for eDNA studies. For example, is it better to sample more sites with less replication or fewer sites with more replication when attempting to characterize nearshore fish communities? Species accumulation curves for fish communities sampled with eDNA suggest that filtering >> 10 L of water is necessary to detect most species present in a system (Gehri et al., 2021; Sard et al., 2019), with an in-depth study of filtering volumes by Cantera et al. (2019) recommending between 34 and 68 L. However, many studies have been able to detect fine-scale heterogeneity using volumes as small as 3 L, filtered in either 3 1-L replicates (Gehri et al., 2021; Kelly et al., 2018; Monuki et al., 2021), or in one 3 L replicate (Port et al., 2016). A goal of our study was to determine whether increasing replication from the more typical 3 1-L replicates to 9 1-L replicates would affect our results. Our power analysis suggested that large effects such as the differences between rock habitat and the other habitats in our study could be identified consistently with lower replication, but that detecting subtle effects such as the habitat-tide interaction likely require additional sampling.

Notably, our study did not investigate potential temporal changes across sampled habitats. Previous eDNA studies have shown that community composition can vary dramatically across relatively short time periods (Beentjes, Speksnijder, Schilthuizen, Hoogeveen, & van der Hoorn, 2019; Berry et al., 2019; Monuki et al., 2021). However, temporal sampling adds logistical considerations, especially increased personnel hours. One technological innovation that can facilitate temporal sampling is an eDNA autosampler, which could be stationed at monitoring locations and take samples at a set frequency (monthly, weekly, daily, hourly, etc.). Multiple efforts are underway to develop affordable autosamplers that could facilitate efficient temporal monitoring in the near future.

For evaluating our laboratory procedure, we investigated the utility of PCR replicates for increasing species detections. Our results are similar to those documented by Beentjes et al. (2019) and Shirazi, Meyer, and Shapiro (2021), who found that PCR replicates were able to identify some new species but that general inferences of community composition did not change with the addition of PCR replicates. However, the PCR replicates in this study received fewer sequencing reads than the primary (“NonRep”) samples (Table S2), and a comparable number of sequencing reads for all replicate samples would provide more conclusive evidence to test the effects of PCR replication. Logistically, PCR replicates increase laboratory costs (reagents, personnel time) and sequencing costs required to achieve sufficient coverage for all samples, but this additional cost may be warranted depending on study goals, especially for detecting rare taxa. In our case collecting additional field samples was relatively straightforward, which should provide more opportunities to sample the dominant fish community at each site.

### Low taxonomic resolution of 12S primer: towards a multi-primer approach

One clear result in our study was that the 12S primer that we used (Riaz et al., 2011) lacked taxonomic resolution for many local species. Despite the 12S primer providing relatively high resolution for many study regions including freshwater lakes in the midwestern USA (Euclide et al., 2021; Gehri et al., 2021; Pukk et al., 2021; Sard et al., 2019), it was unable to provide species-level resolution for some of the most important taxa in Alaska waters including Salmoninae (salmon, also documented by Gehri et al., 2021), Gadidae (cods), Sebastinae (rockfish, also documented by Gold, Sprague, Kushner, Zerecero Marin, & Barber, 2021 with a different 12S marker), and Pleuronectidae (flatfish). Using broadly targeted 12S primers can efficiently assess the presence of many diverse fish taxa with a single metabarcoding primer set; however, primers targeted for specific taxa such as Gadidae or Salmoninae should provide higher resolution and potentially more precise estimates of which species from those families are present and their relative abundance. PCR can lead to deviations between true community composition and eDNA results due to differences in primer efficiencies across species and variation in starting DNA concentrations that can be exacerbated by PCR (Kelly, Shelton, & Gallego, 2019). Therefore, if the primary goal of an eDNA study is to assess important species from multiple taxa, a multi-primer approach with primers targeting specific taxa should produce more accurate results than screening one or a few broadly-targeted primers. This multi-primer approach would require additional effort to design targeted primers, but targeted primers already exist for many of the relevant taxa in southeast Alaska, including salmon (Menning, Simmons, & Talbot, 2020), rockfish (Min, Barber, & Gold, 2021), and eels (Takeuchi et al., 2019).

### Conclusions and recommendations

We used eDNA to investigate nearshore fish communities across habitats at high and low tides in a high-latitude environment and found that communities vary substantially by habitat, with differences between tides specific to certain habitats. We suspect that differences in the fish community across habitats were largely driven by habitat characteristics that influenced how species were detected rather than by differences in species communities themselves. Most notably, we hypothesize that the steep slope of rock habitats led to fewer species detections than more gradually sloping sand and eelgrass habitats. This result provides further evidence of limited eDNA transport in marine systems, with most taxon detections derived from locally abundant species. The effect of tide was more subtle than habitat, with differences in species composition between tides potentially due to movement of small pelagic fish into nearshore in sand habitats at high tide. Additional field samples increased the probability of detecting significant differences across habitats and habitat-tide interactions.

Mounting evidence from our study and many others suggests that marine eDNA reflects diversity at local horizontal and vertical (depth) scales. Future studies could build upon the results presented here by attempting to determine the horizontal or vertical scale over which an eDNA signal dissipates in the nearshore environment, and the extent to which 12S or more targeted species-specific primers might influence the detection range. Further studies could also focus on sampling across tidal stages to detect temporal changes that could be ecologically meaningful. However, our findings indicate that sampling effort is likely better spent by expanding the number of sampling locations rather than sampling across tides.

## Author Contributions

WL, JM, JO, and WD designed the study. Laboratory analysis was conducted by WD. Data analyses were conducted by PB, JM, WL, and DB. All authors assisted in drafting the manuscript and provided feedback and manuscript edits. The scientific results and conclusions, as well as any views or opinions expressed herein, are those of the author(s) and do not necessarily reflect those of NOAA or the Department of Commerce. Any use of trade, firm, or product names is for descriptive purposes only and does not imply endorsement by the U.S. Government.

## Data Archiving Statement

Raw sequence reads (*fastq* format) used in this research along with corresponding metadata are archived with the Dryad Digital Repository (will be uploaded upon acceptance). Scripts used for bioinformatics and taxonomic classifications are available on GitHub (https://github.com/AFSC-Genetics/eDNA_NearshoreMarine).

## Acknowledgements

We thank Katie D’Amelio, Kirby Karpan, and Jackie Whittle for assistance with eDNA sampling and filtering water samples. Funding was provided by a NOAA Office of Habitat Conservation “Refine EFH” grant.

## Figure legends

Fig. S1: Taxon accumulation curves for samples with nine PCR replicates. Replicates are ordered by the one that includes the greatest number of taxa followed sequentially by the replicates that contribute additional unique taxa. The top panel shows the proportion of the total taxa identified cumulatively across the PCR replicates and the bottom panel shows the number of taxa, which varied by sample.

## Tables

Supplementary Table S1: Information on sampling sites including date and time sampled, latitude and longitude, and habitat.

Supplementary Table S2: Read counts by site including negative and positive controls and PCR replicates.

Supplementary Table S3: Number of positive detections for each taxa at each site.

## References

Altschul, S. F., Gish, W., Miller, W., Myers, E. W., & Lipman, D. J. (1990). Basic local alignment search tool. Journal of Molecular Biology, 215(3), 403–410. doi:10.1006/jmbi.1990.9999

Andruszkiewicz, E. A., Koseff, J. R., Fringer, O. B., Ouellette, N. T., Lowe, A. B., Edwards, C. A., & Boehm, A. B. (2019). Modeling environmental DNA transport in the coastal ocean using lagrangian particle tracking. Frontiers in Marine Science, 6(477). doi:10.3389/fmars.2019.00477

Andruszkiewicz, E. A., Starks, H. A., Chavez, F. P., Sassoubre, L. M., Block, B. A., & Boehm, B. (2017). Biomonitoring of marine vertebrates in Monterey Bay using eDNA metabarcoding. PLoS ONE, 12(4), e0176343. doi:10.1371/journal.pone.0176343

Bates, D., Mächler, M., Bolker, B., & Walker, S. (2015). Fitting linear mixed-effects models using lme4. Journal of Statistical Software, 67(1), 1–48. doi:10.18637/jss.v067.i01

Baudron, A. R., Brunel, T., Blanchet, M.-A., Hidalgo, M., Chust, G., Brown, E. J., … Fernandes, P. G. (2020). Changing fish distributions challenge the effective management of European fisheries. Ecography, 43(4), 494–505. doi:https://doi.org/10.1111/ecog.04864

Beaudreau, A. H., Ward, E. J., Brenner, R. E., Shelton, A. O., Watson, J. T., Womack, J. C., … Williams, B. C. (2019). Thirty years of change and the future of Alaskan fisheries: Shifts in fishing participation and diversification in response to environmental, regulatory and economic pressures. Fish and Fisheries, 20(4), 601–619. doi:https://doi.org/10.1111/faf.12364

Beentjes, K. K., Speksnijder, A. G. C. L., Schilthuizen, M., Hoogeveen, M., & van der Hoorn, B. (2019). The effects of spatial and temporal replicate sampling on eDNA metabarcoding. PeerJ, 7, e7335. doi:10.7717/peerj.7335

Berry, T. E., Saunders, B. J., Coghlan, M. L., Stat, M., Jarman, S., Richardson, A. J., … Bunce, M. (2019). Marine environmental DNA biomonitoring reveals seasonal patterns in biodiversity and identifies ecosystem responses to anomalous climatic events. PLoS Genetics, 15(2), e1007943. doi:10.1371/journal.pgen.1007943

Callahan, B. J., McMurdie, P. J., Rosen, M. J., Han, A. W., Johnson, A. J. A., & Holmes, S. P. (2016). DADA2: High-resolution sample inference from Illumina amplicon data. Nature Methods, 13(7), 581–583. doi:10.1038/nmeth.3869

Campbell, N. R., Harmon, S. A., & Narum, S. R. (2015). Genotyping-in-Thousands by sequencing (GT-seq): A cost effective SNP genotyping method based on custom amplicon sequencing. Molecular Ecology Resources, 15(4), 855–867. doi:10.1111/1755-0998.12357

Cantera, I., Cilleros, K., Valentini, A., Cerdan, A., Dejean, T., Iribar, A., … Brosse, S. (2019). Optimizing environmental DNA sampling effort for fish inventories in tropical streams and rivers. Scientific Reports, 9(1), 3085. doi:10.1038/s41598-019-39399-5

Closek, C. J., Santora, J. A., Starks, H. A., Schroeder, I. D., Andruszkiewicz, E. A., Sakuma, K. M., … Boehm, A. B. (2019). Marine vertebrate biodiversity and distribution within the Central California Current using environmental DNA (eDNA) metabarcoding and ecosystem surveys. Frontiers in Marine Science, 6(732). doi:10.3389/fmars.2019.00732

Collins, R. A., Wangensteen, O. S., O’Gorman, E. J., Mariani, S., Sims, D. W., & Genner, M. J. (2018). Persistence of environmental DNA in marine systems. Communications Biology, 1(1), 185. doi:10.1038/s42003-018-0192-6

Craig, P. D., Kellison, G. T., Aaron, J. A., Bronwyn, M. G., Matthew, S. K., Craig, A. L., … Joseph, E. S. (2006). Marine nurseries and effective juvenile habitats: Concepts and applications. Marine Ecology Progress Series, 312, 291-295. Retrieved from https://www.int-res.com/abstracts/meps/v312/p291-295/

Djurhuus, A., Closek, C. J., Kelly, R. P., Pitz, K. J., Michisaki, R. P., Starks, H. A., … Breitbart, M. (2020). Environmental DNA reveals seasonal shifts and potential interactions in a marine community. Nature Communications, 11(1), 254. doi:10.1038/s41467-019-14105-1

Euclide, P. T., Lor, Y., Spear, M. J., Tajjioui, T., Vander Zanden, J., Larson, W. A., & Amberg, J. J. (2021). Environmental DNA metabarcoding as a tool for biodiversity assessment and monitoring: reconstructing established fish communities of north-temperate lakes and rivers. Diversity and Distributions, 27(10), 1966–1980. doi:https://doi.org/10.1111/ddi.13253

Gehri, R. R., Larson, W. A., Gruenthal, K., Sard, N. M., & Shi, Y. (2021). eDNA metabarcoding outperforms traditional fisheries sampling and reveals fine-scale heterogeneity in a temperate freshwater lake. Environmental DNA, 3(5), 912–929. doi:https://doi.org/10.1002/edn3.197

Gold, Z., Sprague, J., Kushner, D. J., Zerecero Marin, E., & Barber, P. H. (2021). eDNA metabarcoding as a biomonitoring tool for marine protected areas. PLoS ONE, 16(2), e0238557. doi:10.1371/journal.pone.0238557

Grüss, A., Pirtle, J. L., Thorson, J. T., Lindeberg, M. R., Neff, A. D., Lewis, S. G., & Essington, T. E. (2021). Modeling nearshore fish habitats using Alaska as a regional case study. Fisheries Research, 238, 105905. doi:https://doi.org/10.1016/j.fishres.2021.105905

Harrison, J. B., Sunday, J. M., & Rogers, S. M. (2019). Predicting the fate of eDNA in the environment and implications for studying biodiversity. Proceedings of the Royal Society B: Biological Sciences, 286(1915), 20191409. doi:doi:10.1098/rspb.2019.1409

Hoaglin, D. C. (1980). A Poissonness plot. The American Statistician, 34(3), 146–149. doi:10.2307/2683871

Hogrefe, K. R., Ward, D. H., Donnelly, T. F., & Dau, N. (2014). Establishing a baseline for regional scale monitoring of eelgrass (Zostera marina) habitat on the lower Alaska Peninsula. Remote Sensing, 6(12), 12447–12477. Retrieved from https://www.mdpi.com/2072-4292/6/12/12447

Holman, L. E., Chng, Y., & Rius, M. (2021). How does eDNA decay affect metabarcoding experiments? Environmental DNA, Online early. doi:https://doi.org/10.1002/edn3.201

Johnson, S. W., Neff, A. D., Thedinga, J. F., Lindeberg, M. R., & Maselko, J. M. (2012). Atlas of nearshore fishes of Alaska: A synthesis of marine surveys from 1998 to 2011. U.S. Dep. Commer., NOAA Tech. Memo. NMFS-AFSC-239, 261 p.

Kelly, R. P., Gallego, R., & Jacobs-Palmer, E. (2018). The effect of tides on nearshore environmental DNA. PeerJ, 6, e4521–e4521. doi:10.7717/peerj.4521

Kelly, R. P., Shelton, A. O., & Gallego, R. (2019). Understanding PCR processes to draw meaningful conclusions from environmental DNA studies. Scientific Reports, 9(1), 12133. doi:10.1038/s41598-019-48546-x

Laurel, B. J., & Rogers, L. A. (2020). Loss of spawning habitat and prerecruits of Pacific cod during a Gulf of Alaska heatwave. Canadian Journal of Fisheries and Aquatic Sciences, 77(4), 644–650. doi:10.1139/cjfas-2019-0238

Legendre, P., & Anderson, M. J. (1999). Distance-based redundancy analysis: Testing multispecies responses in multifactorial ecological experiments. Ecological Monographs, 69(1), 1–24. doi:https://doi.org/10.1890/0012-9615(1999)069[0001:DBRATM]2.0.CO;2

Littlefair, J. E., Hrenchuk, L. E., Blanchfield, P. J., Rennie, M. D., & Cristescu, M. E. 2021). Thermal stratification and fish thermal preference explain vertical eDNA distributions in lakes. Molecular Ecology, 30(13), 3083–3096. doi:https://doi.org/10.1111/mec.15623

Liu, Y., Wikfors, G. H., Rose, J. M., McBride, R. S., Milke, L. M., & Mercaldo-Allen, R. (2019). Application of environmental DNA metabarcoding to spatiotemporal finfish community assessment in a temperate embayment. Frontiers in Marine Science, 6, 674. Retrieved from https://www.frontiersin.org/article/10.3389/fmars.2019.00674

Magoč, T., & Salzberg, S. L. (2011). FLASH: fast length adjustment of short reads to improve genome assemblies. Bioinformatics, 27(21), 2957–2963. doi:10.1093/bioinformatics/btr507

Menning, D., Simmons, T., & Talbot, S. (2020). Using redundant primer sets to detect multiple native Alaskan fish species from environmental DNA. Conservation Genetics Resources, 12(1), 109–123. doi:10.1007/s12686-018-1071-7

Meyer, D., Zeileis, A., & Hornik, K. (2006). The strucplot framework: visualizing multi-way contingency tables with vcd. Journal of Statistical Software, 17(3), 1–48. doi:10.18637/jss.v017.i03

Min, M. A., Barber, P. H., & Gold, Z. (2021). MiSebastes: An eDNA metabarcoding primer set for rockfishes (genus Sebastes). Conservation Genetics Resources. doi:10.1007/s12686-021-01219-2

Monuki, K., Barber, P. H., & Gold, Z. (2021). eDNA captures microhabitat partitioning in a kelp forest ecosystem. bioRxiv, 2021.2006.2001.446542. doi:10.1101/2021.06.01.446542

Newman, D., Berkson, J., & Suatoni, L. (2015). Current methods for setting catch limits for data-limited fish stocks in the United States. Fisheries Research, 164, 86–93. doi:https://doi.org/10.1016/j.fishres.2014.10.018

O’Donnell, J. L., Kelly, R. P., Shelton, A. O., Samhouri, J. F., Lowell, N. C., & Williams, G. D. (2017). Spatial distribution of environmental DNA in a nearshore marine habitat. PeerJ, 5, e3044. doi:10.7717/peerj.3044

Oksanen, J., Blanchet, M. F., Kindt, R., Legendre, P., McGlinn, D., Minchin, P. R., … Wagner, H. (2020). vegan: Community Ecology Package. R package version 2.5-7. https://CRAN.R-project.org/package=vegan.

Port, J. A., O’Donnell, J. L., Romero-Maraccini, O. C., Leary, P. R., Litvin, S. Y., Nickols, K. J., … Kelly, R. P. (2016). Assessing vertebrate biodiversity in a kelp forest ecosystem using environmental DNA. Molecular Ecology, 25(2), 527–541. doi:10.1111/mec.13481

Pukk, L., Kanefsky, J., Heathman, A. L., Weise, E. M., Nathan, L. R., Herbst, S. J., … Robinson, J. D. (2021). eDNA metabarcoding in lakes to quantify influences of landscape features and human activity on aquatic invasive species prevalence and fish community diversity. Diversity and Distributions, 27(10), 2016–2031. Retrieved from https://www.jstor.org/stable/48621937

Riaz, T., Shehzad, W., Viari, A., Pompanon, F., Taberlet, P., & Coissac, E. (2011). ecoPrimers: inference of new DNA barcode markers from whole genome sequence analysis. Nucleic Acids Research, 39(21), e145. doi:10.1093/nar/gkr732

Sard, N. M., Herbst, S. J., Nathan, L., Uhrig, G., Kanefsky, J., Robinson, J. D., & Scribner, K. T. (2019). Comparison of fish detections, community diversity, and relative abundance using environmental DNA metabarcoding and traditional gears. Environmental DNA, 1(4), 368–384. doi:10.1002/edn3.38

Shelton, A. O., Kelly, R. P., O’Donnell, J. L., Park, L., Schwenke, P., Greene, C., … Beamer, E. M. (2019). Environmental DNA provides quantitative estimates of a threatened salmon species. Biological Conservation, 237, 383–391. doi:https://doi.org/10.1016/j.biocon.2019.07.003

Shirazi, S., Meyer, R., & Shapiro, B. (2021). Revisiting the effect of PCR replication and sequencing depth on biodiversity metrics in environmental DNA metabarcoding. Authorea. doi:10.22541/au.159309876.62184178/v2

Sigsgaard, E. E., Torquato, F., Frøslev, T. G., Moore, A. B. M., Sørensen, J. M., Range, P., … Thomsen, P. F. (2020). Using vertebrate environmental DNA from seawater in biomonitoring of marine habitats. Conservation Biology, 34(3), 697–710. doi:10.1111/cobi.13437

Skern-Mauritzen, M., Olsen, E., & Huse, G. (2018). Opportunities for advancing ecosystem-based management in a rapidly changing, high latitude ecosystem. Ices Journal of Marine Science, 75(7), 2425–2433. doi:10.1093/icesjms/fsy150

Steele, M. A., Schroeter, S. C., & Pace, H. M. (2006). Experimental evaluation of biases associated with sampling estuarine fishes with seines. Estuaries and Coasts, 29(6), 1172–1184. doi:10.1007/BF02781818

Stoeckle, M. Y., Adolf, J., Charlop-Powers, Z., Dunton, K. J., Hinks, G., & VanMorter, S. M. (2020). Trawl and eDNA assessment of marine fish diversity, seasonality, and relative abundance in coastal New Jersey, USA Ices Journal of Marine Science, 78(1), 293–304. doi:10.1093/icesjms/fsaa225

Takeuchi, A., Sado, T., Gotoh, R. O., Watanabe, S., Tsukamoto, K., & Miya, M. (2019). New PCR primers for metabarcoding environmental DNA from freshwater eels, genus Anguilla. Scientific Reports, 9(1), 7977. doi:10.1038/s41598-019-44402-0

Thomsen, P. F., Kielgast, J., Iversen, L. L., Møller, P. R., Rasmussen, M., & Willerslev, E. (2012). Detection of a diverse marine fish fauna using environmental DNA from seawater samples. PLoS ONE, 7(8), e41732. doi:10.1371/journal.pone.0041732

Thorson, J. T., & Barnett, L. A. K. (2017). Comparing estimates of abundance trends and distribution shifts using single- and multispecies models of fishes and biogenic habitat. Ices Journal of Marine Science, 74(5), 1311–1321. doi:10.1093/icesjms/fsw193

Weingartner, T., Eisner, L., Eckert, G. L., & Danielson, S. (2009). Southeast Alaska: oceanographic habitats and linkages. Journal of Biogeography, 36(3), 387–400. doi:https://doi.org/10.1111/j.1365-2699.2008.01994.x

West, K. M., Stat, M., Harvey, E. S., Skepper, C. L., DiBattista, J. D., Richards, Z. T., … Bunce, M. (2020). eDNA metabarcoding survey reveals fine-scale coral reef community variation across a remote, tropical island ecosystem. Molecular Ecology, 29(6), 1069–1086. doi:https://doi.org/10.1111/mec.15382

