## Supplementary figures and images for "Leveraging eDNA metabarcoding to characterize nearshore fish communities in Southeast Alaska: Do habitat and tide matter?"

### Supplemental Figure S1

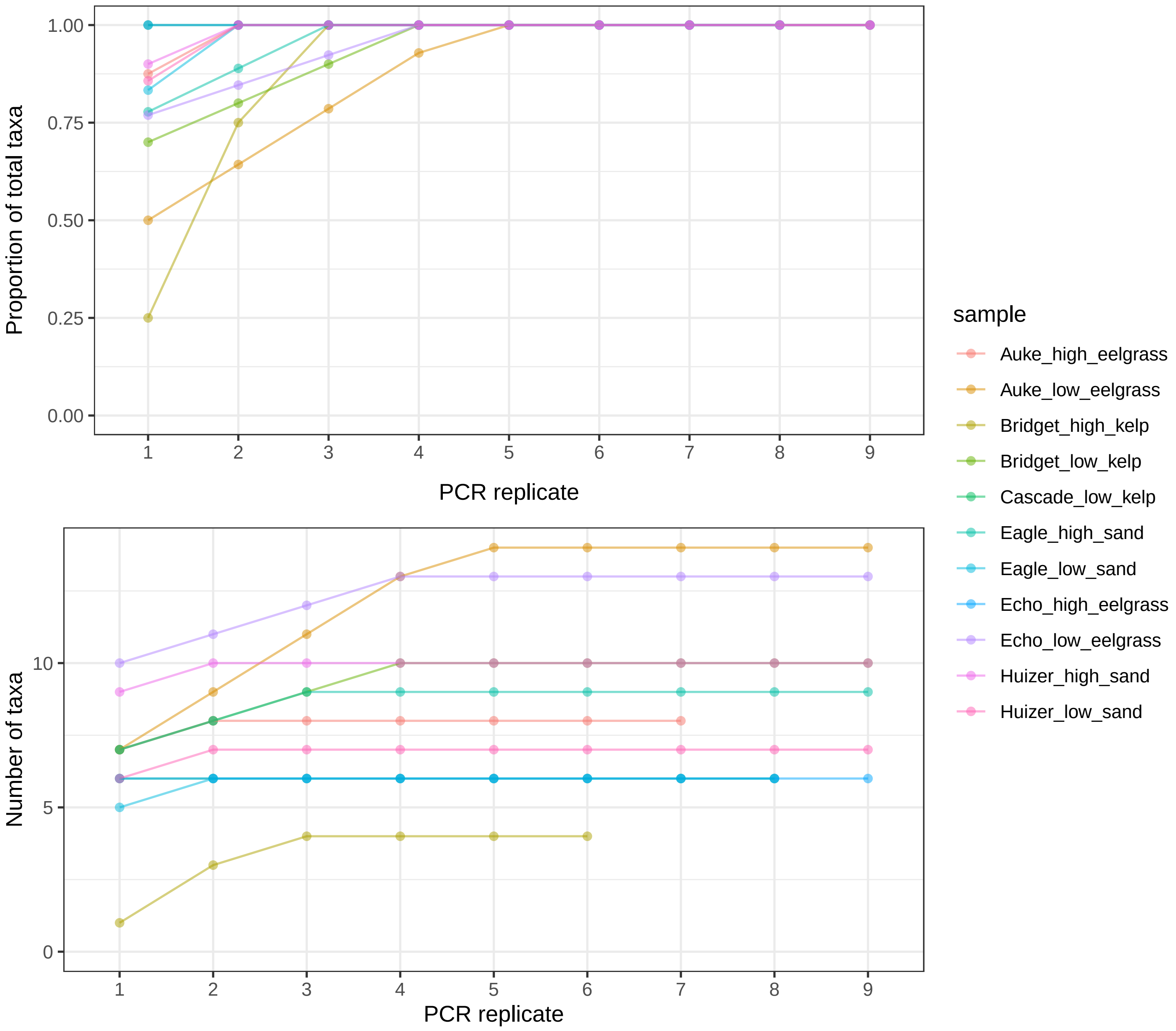
